# Reward-related regions form a preferentially coupled system at rest

**DOI:** 10.1101/304865

**Authors:** Jeremy F. Huckins, Babatunde Adeyemo, Fran M. Miezin, Jonathan D. Power, Evan M. Gordon, Timothy O. Laumann, Todd F. Heatherton, Steven E. Petersen, William M. Kelley

## Abstract

Neuroimaging studies have implicated a set of striatal and orbitofrontal cortex (OFC) regions that are commonly activated during reward processing tasks. Resting-state functional connectivity (RSFC) studies have demonstrated that the human brain is organized into several functional systems that show strong temporal coherence even in the absence of goal-directed tasks. Here we use seed-based and graph-theory RSFC approaches to characterize the systems-level organization of putative reward regions of at rest. Peaks of connectivity from seed-based RSFC patterns for the nucleus accumbens (NAcc, anatomically defined) and orbitofrontal cortex (OFC, functionally defined) were used to identify candidate reward regions which were merged with a previously used set of regions (Power et al., 2011). Graph-theory was then used to determine system-level membership for all regions. Several regions previously implicated in reward-processing (NAcc, lateral and medial OFC, and ventromedial prefrontal cortex) comprised a distinct, preferentially coupled system. This RSFC system is stable across a range of connectivity thresholds and shares strong overlap with meta-analyses of task-based reward studies. This reward system shares between-system connectivity with systems implicated in cognitive control and self-regulation, including the fronto-parietal, cingulo-opercular, and default systems. Further, differences may exist in the pathways through which control systems interact with key regions of this reward system. Whereas NAcc regions of the reward system are functionally connected to cingulo-opercular and default systems, OFC regions of the reward system show stronger connectivity with the fronto-parietal system. We propose that future work may be able to interrogate group or individual differences in connectivity profiles using the regions delineated in the current work to explore potential relationships to appetitive behaviors, self-regulation failure, and addiction.

## Introduction

Reward is a multifaceted term which captures multiple components of cognitive experience and behavior related to liking, wanting and learning (Berridge & Kringelbach, 2008). Non-human animal studies show that consuming rewards (foods, drugs) or engaging in rewarding activities (i.e., sex) is associated with activation of mesolimbic dopamine brain regions (e.g., the ventral tegmental area and nucleus accumbens/ventral striatum) and the orbitofrontal cortex (Boileau et al., 2003; Carelli, Ijames, & Crumling, 2000; Damsma, Pfaus, Wenkstern, Phillips, & Fibiger, 1992; Everitt, 1990; M. L. Kringelbach, 2005; Schilström, Svensson, Svensson, & Nomikos, 1998). In humans, functional neuroimaging work has similarly shown that activity in the ventral striatum and orbitofrontal cortex (OFC) increases during both reward consumption (Breiter et al., 1997; Gottfried, O’Doherty, & Dolan, 2003; Kringelbach, O’Doherty, Rolls, & Andrews, 2003) and reward anticipation, such as when viewing a food advertisement or when seeing or smelling a cigarette (Carter & Tiffany, 1999; Harris, Bargh, & Brownell, 2009; Lambert, Neal, Noyes, Parker, & Worrel, 1991; Rapuano, Huckins, Sargent, Heatherton, & Kelley, 2016; Sayette & Hufford, 1997). In addition to activation of the ventral striatum, and OFC, human neuroimaging work has consistently shown reward-related activity extending into the thalamus, the dorsal striatum, VTA, medial prefrontal cortex, and insula in response to reward cues such as food images, drugs, attractive faces, as well as to secondary reward cues such as money (Cloutier, Heatherton, Whalen, & Kelley, 2008; Due, Huettel, Hall, & Rubin, 2002; Garavan et al., 2000; Knutson, Taylor, Kaufman, Peterson, & Glover, 2005; Sescousse, Caldú, Segura, & Dreher, 2013; Somerville, Hare, & Casey, 2011; van der Laan, de Ridder, Viergever, & Smeets, 2011; Yarkoni, Poldrack, Nichols, Van Essen, & Wager, 2011).

More recently, the field of neuroscience has demonstrated a relationship between reward cue-reactivity and real-world appetitive cravings and behaviors such as weight gain, and sexual interest (Demos, Heatherton, & Kelley, 2012; Janes et al., 2010; Lopez, Hofmann, Wagner, Kelley, & Heatherton, 2014; McClernon, Kozink, & Rose, 2008; Stice, Yokum, Bohon, Marti, & Smolen, 2010). Importantly, experimentally-measured reward responsivity can reflect well-established sensitivities to rewards. Casey and colleagues (2011) showed that individuals who had difficulty delaying gratification as a child (e.g., Mischel et al., 1989) exhibited heightened reward responsivity in the ventral striatum when viewing appetitive cues over 40 years later. Given the common pattern of activation across task-based studies of reward and their relation to individual differences in reward motivation and behavior, researchers have posited a putative human “reward system” that functions to represent reward signal strength and to motivate subsequent behavior towards reward cues (Koob & Volkow, 2010; Volkow, Wang, Fowler, Tomasi, & Telang, 2011).

Converging evidence in support of this idea comes from studies examining resting-state functional connectivity (RSFC) patterns between the ventral striatum and other brain regions (Barnes et al., 2010; Cauda et al., 2011; Choi, Yeo, & Buckner, 2012; Di Martino et al., 2008). RSFC measures the degree to which spontaneous activity across brain regions correlates at rest (i.e., in the absence of explicit task-constraints) (Biswal et al., 1995). RSFC signal correlations are believed to reflect histories of co-activation across brain regions—a pattern of statistical coherence that arises throughout development and provides a measure of the long-term functional relatedness of brain regions (Crossley et al., 2013; Dosenbach et al., 2010). Thus far, the majority of neuroimaging studies examining RSFC in putative reward regions have primarily adopted *seed-based* approaches whereby a seed is placed in one brain region and the analysis identifies other brain regions with similar spontaneous fluctuations in activity. Such seed-based studies of ventral striatum RSFC have commonly identified correlated activity in orbitofrontal and ventral regions of prefrontal cortex, posterior cingulate cortex, inferior parietal lobule, thalamus, hippocampus and caudate (Barnes et al., 2010; Cauda et al., 2011; Choi et al., 2012; Di Martino et al., 2008). Additonally, Choi and colleagues (2012) determined that the ventral striatum was primarily connected to a “limbic” system which included ventral prefrontal cortex in their 17 system parcellation. There seems to be considerable correspondence between the extant seed-based RSFC maps and the task-based maps, lending credence to the notion that putative reward regions are preferentially activated and coupled, across both task and resting-state studies.

The notion of a reward system makes intuitive sense given the converging evidence from meta-analyses of task-based reward studies and seed-based RSFC studies; however, an open question is whether the reward-related activations that are commonly observed across task-based studies of reward and in seed-based maps of RSFC constitute a discrete functional system, in which reward-related regions are preferentially connected to each other at rest, or whether reward cue-reactivity tasks engage brain regions across multiple functional systems. We know that a wide-range of regions process reward-related information, is there a distinct subset of regions which are preferentially connected to each other at rest which include regions typically thought to part of the human reward system. Put simply, is there a set of brain regions that constitute a preferentially coupled functional reward system in the human brain at rest?

One fruitful way to identify and characterize discrete brain systems is to apply network analyses to RSFC data. In particular, graph-theory-based network analysis is a powerful approach to examining RSFC that reveals the integrity of brain systems as well as how connectivity between systems function within wider network contexts (Power et al., 2011). Within the context of graph-theory-based network analysis of RSFC data, there are a variety of concepts which are critical to fully understand before proceeding. When observing a RSFC network, there are nodes (brain regions) which are connected by ties (correlation between those regions). When observing a network, if you included all of the connections across 264 regions (say from Power et al., 2011) you would have to account for over 34,000 connections. One common method with which many researchers chose to reduce the number of connections they are observing, is to look at only the strongest X percentage of connections, which is often called tie-density. A tie-density of 4% means that the highest 4% of connections are retained and the rest of the connections are excluded. If one were to choose one specific tie-density, then it would be a relatively arbitrary thresholding of the network, and could identify properties or characteristics which are not representative of the network as a whole. To reduce the reliance of results on a single tie-density, it is preferred to look for consistent results across a variety of tie-densities. At low tie-densities, which include a relatively low % of connections (<5%) a highly segmented network could be observed, with many subcomponents. At higher tie-densities (<10%) many of these subcomponents would merge, leads to a highly connected network with few subcomponents. Above a tie-density of 20% a network with 200+ nodes will generally look highly connected mush with few distinguishable subcomponents.

To observe sparse (only selecting certain connections) networks such you could map it out in an anatomical manner, or alternatively, it can be mapped by connectivity strength so that subcomponents of the network which are strongly coupled are near each other. An example of this is spring-embedded graphing, a method where springs between nodes where stronger connectivity is mapped as a stronger spring between the nodes, pulling them together while weaker connectivity is represented as a weaker spring between the nodes, or for connections below the tie-density threshold, no spring is used. This representation helps position the nodes such that groups of nodes which are more strongly coupled are closer together and those that are less strongly coupled are further apart, giving a functional representation of the network. An important method which can greatly inform our understanding of network components and structure is community assignments. There are a variety of methods, including Infomap, a random walk algorithm, which can identify where systems or subnetworks are located.

Identifying systems which are present across a wide-variety of tie-densities would suggest that they are relatively robust within the set of nodes being tested. Furthermore, permutation testing by selecting different sets of individuals from a cohort then performing the community assignment on that subset of individuals allows for formal testing of co-occurance of regions in a community. Less formally, this allows us to ask if regions in a system are preferentially connected to each other in a reliable quantitative manner.

While tie-density is often used as a method to identify important connections but selecting the ones with the strongest temporal correlation, multiscale backbone is a method which tried to identify relevant connections across a variety of thresholds. An example where this might be relevant is looking at the network of air travel across the world. Using tie-density to select connections would identify only the busiest connections between major airports such as New York City’s JFK and London’s Heathrow while excluding all connections to smaller, local airports such as Lebanon Municipal Airport in New Hampshire. Tie-density is very useful for thresholding the network so it can be submitted to community detection methods. One weakness of tie-density is that its less accurate at recapitualiting the entire network. A model which identified important connections at local levels, important connections at a national level and important connections at a global level might be able to do a better job recapitulating the whole network when combined. This type of method is multiscale backbone analysis of a network. In the current example, multiscale backbone analysis would be particularly useful for identifying how to get from airport A to airport B where tie-density would only be useful for this if the two airports you cared about were major global hubs. In applying this example to brain data we can identify the multiscale backbone which could be thought of as a identifying connections across multiple levels that could transfer information. While we are not able to directly test this information transfer property using resting-state it does give us insight into possible paths through which information could propagate between systems which is particularly relevant within the context of reward and self-regulation of reward.

Network-based approaches to understanding brain organization and function (Power et al., 2011) have already identified resting-state systems corresponding to sensory processing, motor control, attention, and, importantly, additional resting-state systems that likely support aspects of self-regulation—the fronto-parietal system, the cingulo-opercular system, and the default mode system (Corbetta, 1998; Dosenbach et al., 2006; Shulman et al., 1997) that may function to regulate putative reward-related activity (Kelley et al., 2015; Lopez et al., 2017; Somerville et al., 2011; Weiland et al., 2013; Zanto & Gazzaley, 2013). Several regions within the default mode network including regions of the medial prefrontal cortex and poster cingulate cortex have been linked to the representation of self (Kelley et al., 2002; Moran, Macrae, Heatherton, Wyland, & Kelley, 2006) and these regions may play an important role in establishing long-term goals consistent with one’s sense of self. The cingulo-opercular system has been described as a core system for the maintenance and monitoring of long term goals and task sets (Dosenbach et al., 2007; Sadaghiani & D’Esposito, 2015). Activity within three key regions of this system— the left and right anterior insula/frontal operculum and the dorsal anterior cingulate extending into the middle superior frontal cortex all show a robust, domain general spike in activity at the beginning of goal-directed task performance that is maintained in a tonic manner throughout the task and is sensitive to performance related feedback. The fronto-parietal system incorporates dorsolateral PFC regions, posterior parietal and inferotemporal regions that are commonly activated during tasks that place demands on inhibitory control, working memory, and attentional filtering (Dosenbach et al., 2007). Together, these systems contain many of the cortical regions which are typically identified in event-related fMRI studies of self-regulation.

Reproducibility of functional architechture and systems across individuals and laboratories suggests that RSFC has a common architecture (Biswal et al., 2010). Noticeably absent from network-based RSFC analyses thus far, however is evidence of a distinct reward system that resembles the sets of brain regions that are commonly active during task-based studies of reward. This is despite robust overlap in activation patterns between task-based studies of reward and *seed-based* RSFC studies of key reward regions such as the ventral striatum. One possibility is that the brain regions showing strong seed-based functional connectivity with the ventral striatum are pieces of separate systems that are jointly recruited during tasks that promote reward processing. If so, then the notion of a true “reward system” may be misleading. Alternatively, prior network-based studies of RSFC failed to identify a reward system in part because of the way nodes are chosen for inclusion in graph-theory analyses. For example, Power and colleagues (2011) used meta-analyses of task-based studies to define a set of spherical nodes distributed throughtout the brain for further analysis. This approach provided robust, but not full, brain coverage, omitting nodes for the ventral striatum and other subcortical areas which are difficult to define using large spherical nodes. In short, community assignments in network-based RSFC analyses can be highly dependent on how brain nodes are chosen, defined, and thresholded whereas seed-based RSFC approaches do not readily permit ways to assess community assignments. Collectively, however, seed-based RSFC and graph-theory network analyses may be complementary— seed-based approaches may be useful in (1) identifying potential target regions which can then (2) be formally tested for community membership using graph theoretical analyses.

The current work capitalizes on both RSFC approaches in a large cohort of healthy individuals. We used seed-based RSFC patterns from putative reward regions (i.e., the ventral striatum and lateral OFC) to identify *candidate* reward system regions with anatomically defined NAcc and functionally defined OFC seeds. We extended previous work by Power and colleagues (2011) using graph theoretical techniques to determine if these regions comprised a distinct reward system in the human brain. Three possible outcomes were considered: 1) no resting-state reward system (i.e., all of the candidate nodes are members of other RSFC systems); 2) all of the candidate reward regions clustered into a distinct system, and 3) a subset of the regions comprised a distinct system. Specifically, we hypothesize that reward-specific regions (critically NAcc and OFC) will be preferentially clustered into a distinct community, following the third option above. Characterizing where the communities that candidate reward regions were members of and how they interact may prove critical in understanding individual differences that permit some individuals to successfully resist unhealthy temptations in daily life and lead others to self-regulation failures.

## Material and Methods

### Subjects

Subjects were 1,016 individuals from the Dartmouth College community. The results presented here are from 828 subjects (580 females) with a mean age of 20.8 +/- 3.7 years old (range: 18-49), who passed stringent RSFC screening measures developed by Power et al. (2014), described below. Subjects had normal or corrected-to-normal visual acuity. Each subject provided informed consent in accordance with the guidelines set by the Committee for the Protection of Human Subjects at Dartmouth College and received either course credit or monetary compensation for participating in the study.

### Apparatus

Imaging was performed on a Philips Intera Achieva 3-Tesla scanner (Philips Medical Systems, Bothell, WA) using a 32-channel phased array head coil. During scanning, participants viewed a white fixation cross on a black background projected on a screen positioned at the head end of the scanner bore, which participants viewed through a mirror mounted on top of the head coil.

### Imaging

Anatomic images were acquired using a high-resolution 3-D magnetization-prepared rapid gradient echo sequence (MP-RAGE; 160 sagittal slices; TE, 4.6 ms; TR, 9.9 ms; flip angle, 8°; voxel size, 1 × 1 × 1 mm). Resting-state functional images were collected using T2*-weighted fast field echo, echo planar functional imaging sensitive to BOLD contrast (TR= 2500 ms; TE= 35 ms; flip angle= 90°; 3 × 3 mm in-plane resolution; sense factor of 2). Functional scanning was performed in two runs; during each run, 120 or 240 axial images (36 slices, 3.5 mm slice thickness, 0.5 mm skip between slices) were acquired, allowing complete brain coverage. As such, each participant completed between 10 and 20 minutes of RSFC scanning as data was collected as part of multiple experiments.

### RSFC Analyses

All processing was performed exactly as in Power et al. (2014) with two exceptions: frame-displacement (FD) threshold was set to 0.25mm (instead of 0.2mm) and 36 motion parameters (instead of 24) were used for motion regression. Functional images were preprocessed to reduce artifacts, including: (i) slice-timing correction, (ii) rigid body realignment to correct for head movement within and across runs, (iii) within-run intensity normalization such that the intensity of all voxels and volumes achieved a mode value of 1000 scale with 10 units equal to ~1% signal change, (iv) transformation to a standardized atlas space (3 mm isotropic voxels) based on (Talairach & Tournoux, 1988), (v) frame censoring, (vi) demeaning and detrending each functional run, (vii) nuisance regression (excluding censored frames), (viii) interpolation, (ix) bandpass filtering (0.009 < f < 0.08Hz) and (x) spatial blurring using a 6mm FWHM kernel following Power et al. (2014). Final correlation calculations between time-courses were calculated based upon *uncensored* frames. Preprocessing steps i-v were completed using custom scripts which call 4dfp Tools (ftp://imaging.wustl.edu/pub/raichlab/4dfp_tools/). Steps specific to resting-state function-connectivity processing (vi-x) were completed using custom MATLAB (Version R2012b, by MathWorks, Natick, MA) scripts.

### Nuisance regressors

To control for motion, a Volterra expansion (Friston, Williams, Howard, Frackowiak, & Turner, 1996) with 36 motion parameters was used. This expansion includes motion (R), motion squared (R^2^), motion at the previous frame (R_t-1_) and motion in the previous frame squared (R_t-1_^2^). Tissue-based nuisance regressors were calculated by taking the mean signal across voxels within each of the following individual masks from FreeSurfer (http://surfer.nmr.mgh.harvard.edu) (Dale, Fischl, & Sereno, 1999; Desikan et al., 2006): an eroded (up to 4x) ventricular mask for the cerebro-spinal fluid, an eroded white matter mask for the white matter signal, and a whole-brain mask for global signal. When eroded masks included no voxels, lesser erosions were progressive considered until a mask with qualifying voxels was identified. This occurred infrequently for white-matter masks while erosions of 1 were often used for CSF masks. The first derivative for each tissue regressor, as calculated by the difference from the current from to the previous frame, was also included.

### Volume censoring and data retention

Movement of the head from one volume to the next (FD) was calculated by the sum of the absolute values of the differentiated realignment values (x, y, z, pitch, roll, yaw) at each time-point (Power, Barnes, Snyder, Schlaggar, & Petersen, 2012). A frame displacement threshold of 0.25mm was used. Volumes with motion above the frame displacement threshold were identified and replaced after multiple regressions but prior to frequency filtering. Spectral decomposition of the uncensored data was performed and used to reconstitute (stage vii: interpolation) data at censored time-points. The frequency content of uncensored data was calculated with a least squares spectral analyses for non-uniformly sampled data (Mathias et al., 2004) based upon the Lomb-Scargle periodogram (Lomb, 1976). Segments of data with less than 5 contiguous volumes below the FD threshold were flagged for censoring. Functional runs were only included in the final analysis if the run contained 50 or more uncensored frames. Only subjects with at least 120 frames of uncensored data across runs were included in the current study (N=828). Consistent with Power et al., (2014), only uncensored volumes were used when calculating temporal correlations.

Using these metrics, 828 of 1,016 individuals passed RSFC processing with more than 120 uncensored frames. Temporal masks retained 79% +/- 14% (range: 26%-98%) of the data across the 828 included subjects. Subjects retained, on average, 238.5 +/- 93.9 frames (range: 121 – 470).

### Reward regions of interest

Regions-of-interest (ROIs) for the left and right NAcc were individually defined for each subject based on an automated segmentation of the high-resolution MPRAGE anatomical image using FreeSurfer’s automated parcellation tool aseg (Fischl et al., 2004). Voxel-based RSFC maps were then generated for each subject using their individually-defined nucleus accumbens (NAcc) ROIs. Bilateral OFC regions were defined based peaks of NAcc connectivity, which corresponded to peaks identified in prior task-based studies of reward (Wagner, Boswell, Kelley, & Heatherton, 2012) conducted in a subset of the same subjects included in the RSFC analyses reported here. These peak OFC regions shared strong overlap with OFC peaks identified in the task-based meta-analysis of reward (Yarkoni et al., 2011).

### Neurosynth analysis

To determine the extent to which RSFC was similar to *task-based* activation patterns observed in studies of reward, RSFC seed maps were compared to an automated meta-analysis performed using Neurosynth (Yarkoni et al., 2011). Mean bilateral NAcc and OFC RSFC seed maps for all 828 subjects were submitted to the Neurosynth Image Decoder which calculates the correlation between an activation map and a statistical image for each of the 3000+ feature terms in the Neurosynth database (Table I).

**Table I.**
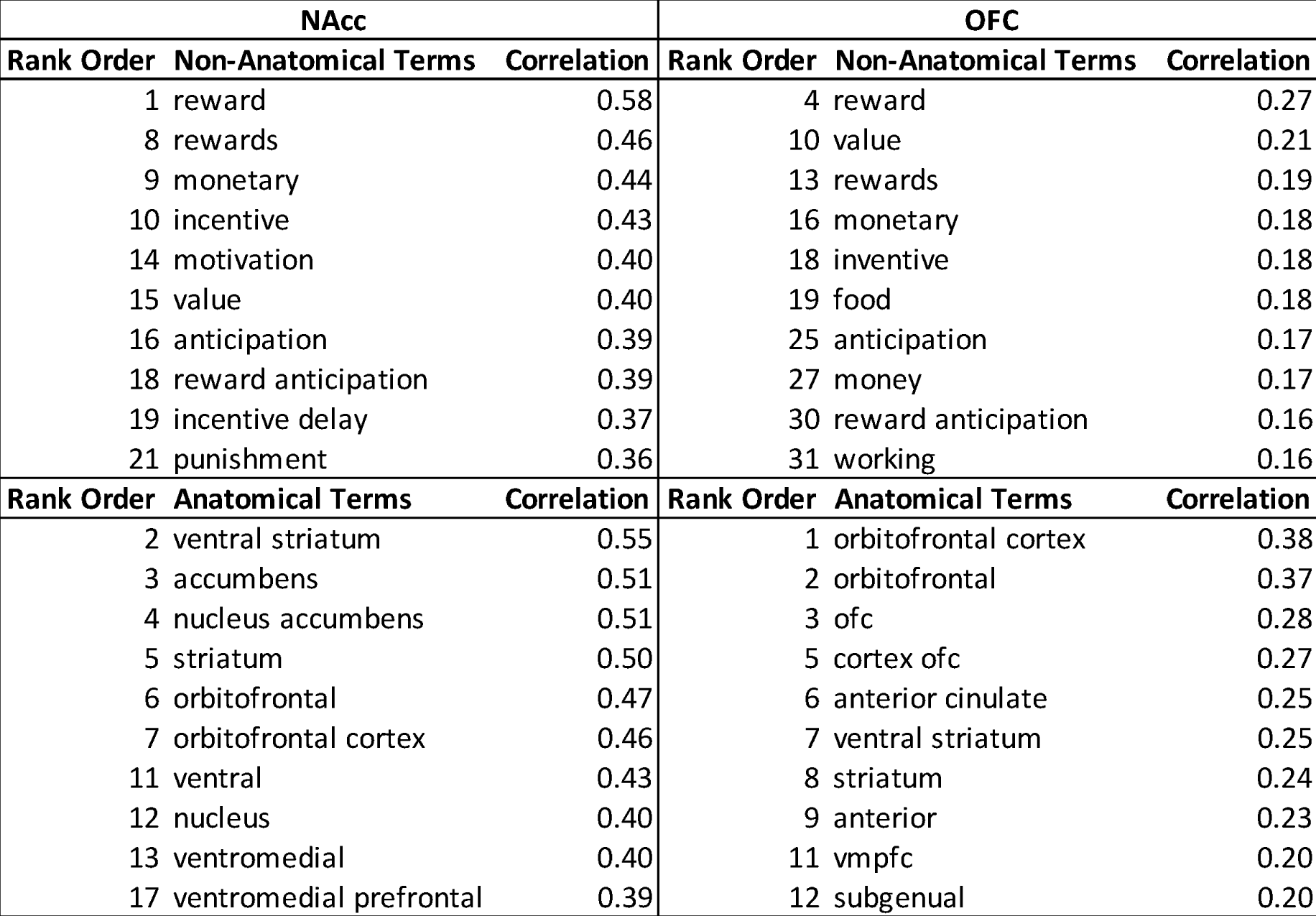
Top 10 non-anatomical and anatomical feature term correlations between seed-based NAcc (left) and OFC (right) RSFC and task-based reverse-inference Z-score images from the Neurosynth Decoder (Yarkoni et al., 2011).

### Graph analysis

Graph theoretical community assignments of RSFC data have been performed on regions previously identified through task-based meta-analyses (Power et al., 2011), automated whole-brain anatomical parcellation (Davis et al., 2013) and RSFC-boundary mapping (Gordon et al., 2014). Community assignment algorithms can then be used to identify systems (i.e., sub-networks) within the graph. Community assignments derived from approaches like these are dependent on how nodes are defined for the overall network and the tie-density (connectivity threshold retaining the strongest X percentage of connections) range of values considered.

Although prior graph theoretical community assignments have been performed on a network of nodes identified through meta-analyses of task-based studies (e.g., Power et al., 2011), none of the currently available parcellation schemes were optimized to include nodes from task-based studies of reward. Indeed, the initial parcellation scheme employed by Power et al. (2011) did not include subcortical NAcc nodes.

In the present study, we modified the existing set of regions (N = 264, 10 mm spheres) from Power et al. (2011) to include peaks identified from the NAcc and OFC RSFC group (N = 828) seed maps. NAcc seeds were derived from individual anatomical segmentation, while OFC seeds were based on group-map peaks of NAcc RSFC connectivity (MNI: 23, 33, −13; −24, 35, −13), which also showed strong similarity to regions identified in task-based reward literature (Wagner et al., 2012). Nodes modified from the original 264 for graph analysis include left, right and medial OFC [BA 10/11], left and right dorsal frontal cortex [BA 46], left lateral prefrontal cortex [BA 9], left inferior medial temporal gyrus [BA 37], left inferior parietal lobule [BA 40], ventral tegmental area (VTA), bilateral thalamus, hippocampus, caudate and right putamen. Where possible, subcortical regions were defined anatomically on a subject-by-subject basis using FreeSurfer auto-segmentation (left and right NAcc, left and right caudate, right putamen, left and right thalamus, and left and right hippocampus). Cortical regions that were added or modified were added as 10 mm spheres to maintain consistency with Power et al., 2011. The VTA was defined as a 4mm spherical seed. Nodes from Power et al., 2011 within 20 mm of the center of any regions added to the new dataset were excluded from the updated regions set. These modifications resulted in a final set of 271 regions, with modifications to Power and colleagues (2011) described in Table II.

**Table II.**
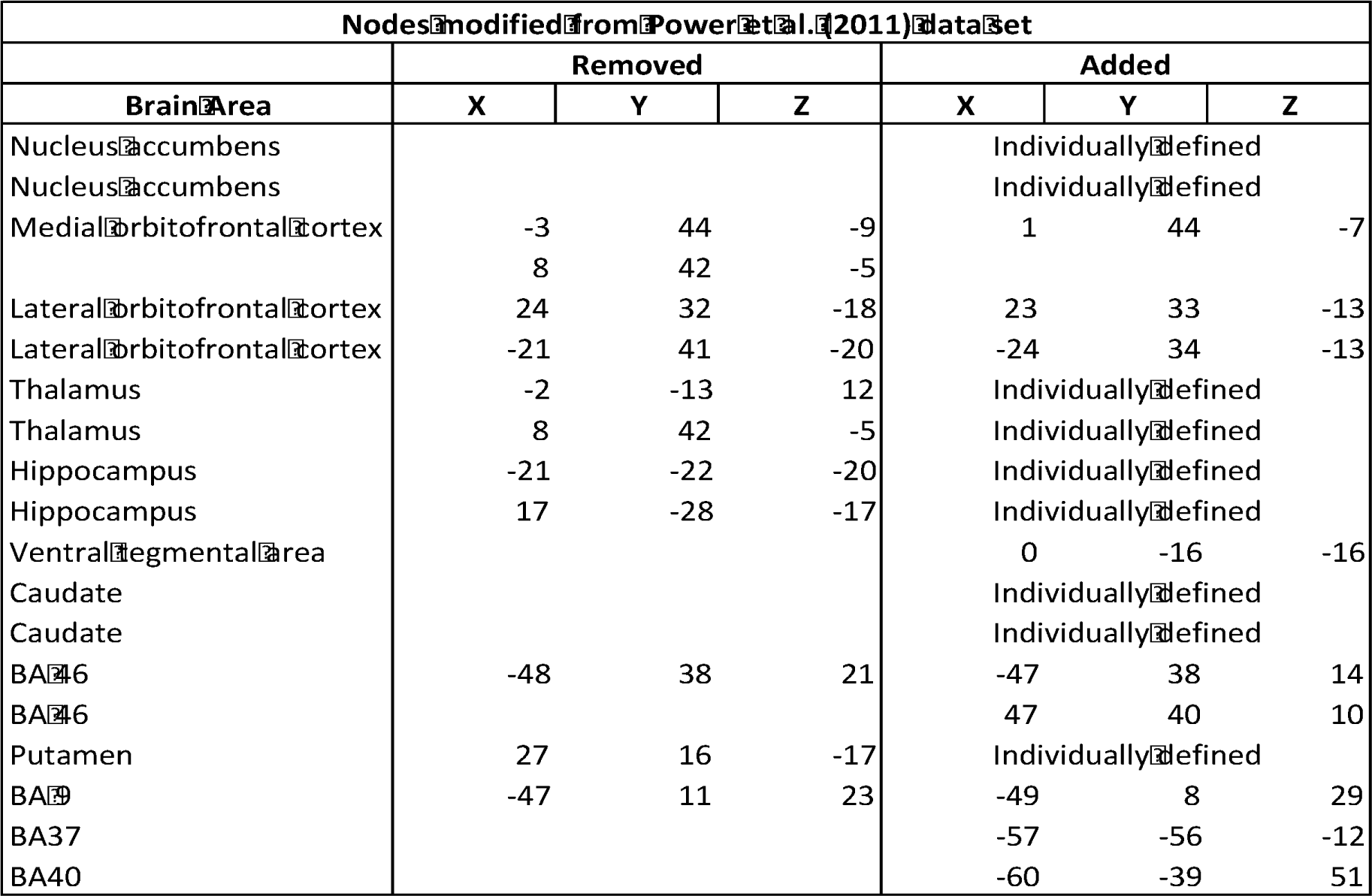
Peak coordinates are listed for all supplemental nodes and the nodes they replaced from the Power et al. (2011) parcellation scheme.

Average time-courses for each node were calculated and cross-correlated with the time-courses from every other node to create a 271 × 271 correlation matrix for each subject. Mean connectivity matrices were then created and transformed into thresholded graphs across a wide range of tie-densities (strongest 2% to 20% of connections retained; Power et al., 2011). Network structure was assessed using the Infomap algorithm, optimal solution to the map equation, a random-walk community assignment method (Rosvall & Bergstrom, 2008) as implemented in Graphtools 1.0 available from www.nitrc.org/projects/graphtools/. Permutation testing to assess stability of the networks in a quantitative manner was done by performing 1000 permutations, with each permutation randomly selecting 25 individuals. Infomap clustering was performed on each permutation and the pairwise co-occurrence of regions within a cluster was recorded. Regions are considered to be significantly coupled if they are in the same community 95% or more of the time and form an independent community, rejecting the null-hypothesis that they are not clustered.

The network was visualized using a spring-embedded graph (Figure 2, bottom), a type of force directed graph drawing, where all ties which are present at the given tie-density, are included as springs between nodes where stronger connectivity is mapped as a stronger spring between the nodes, pulling them together while weaker connectivity is represented as a weaker spring between the nodes, or for connections below the tie-density threshold, no spring is used. This representation helps position the nodes such that groups of nodes which are more strongly coupled are closer together and those that are less strongly coupled are further apart, giving a functional representation of the network. Community assignments from Infomap were used to color nodes in the spring-embedded graph.

**Figure 1.**
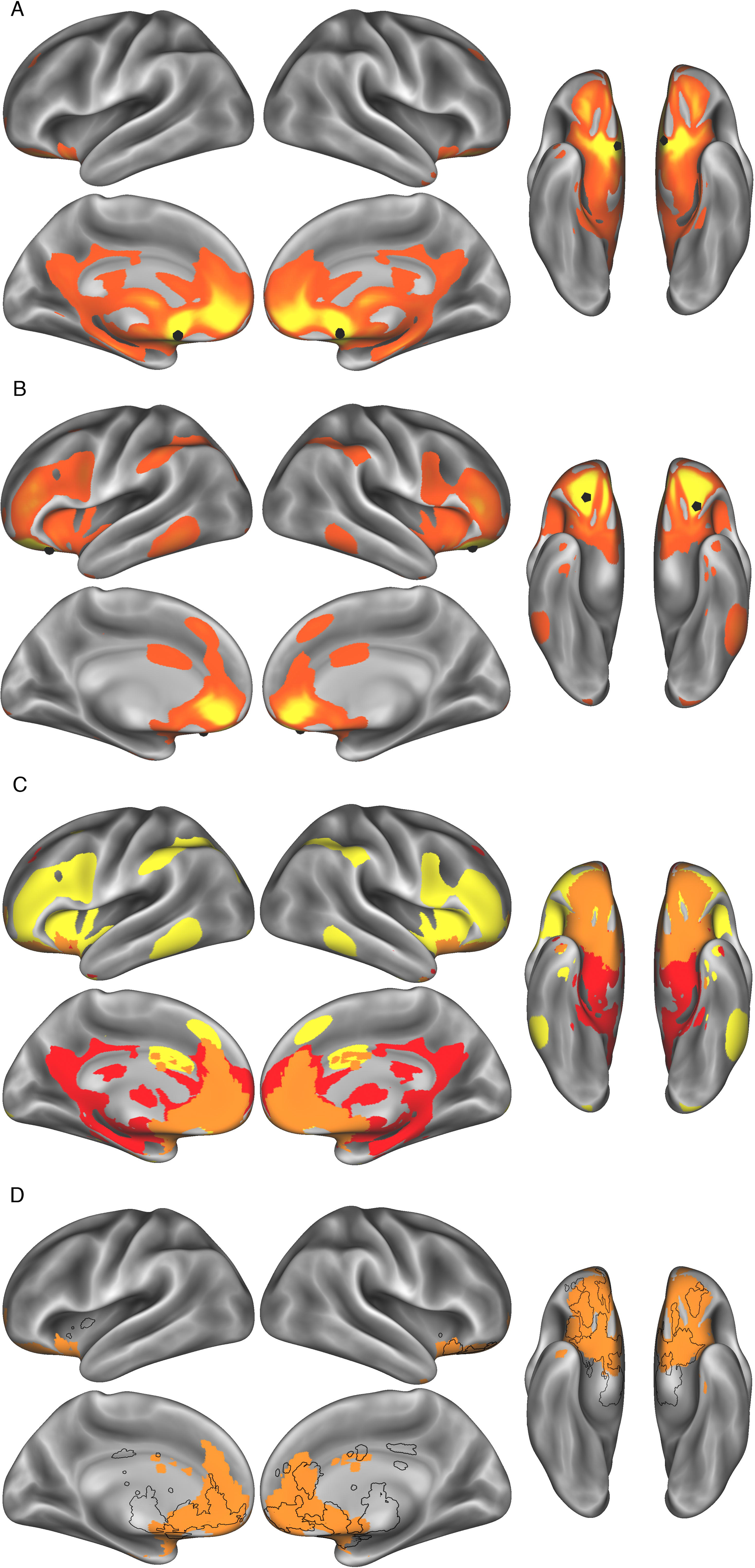
Seed-based RSFC for NAcc (A) and OFC (B). Independent connectivity for NAcc (C, red) and OFC (C, yellow) are displayed on the same surface, shown with shared connectivity (C, orange). Brain regions that are commonly activated in a meta-analysis of task-based studies on the feature term "reward" (D) (Yarkoni et al., 2011) are shown in black outline highlights over shared connectivity of the NAcc and OFC, shown in orange. RSFC data are thresholded at r > 0.05 and are displayed on inflated lateral, medial, and ventral cortical surfaces. Seed locations for NAcc and OFC are highlighted as black spheres.

**Figure 2.**
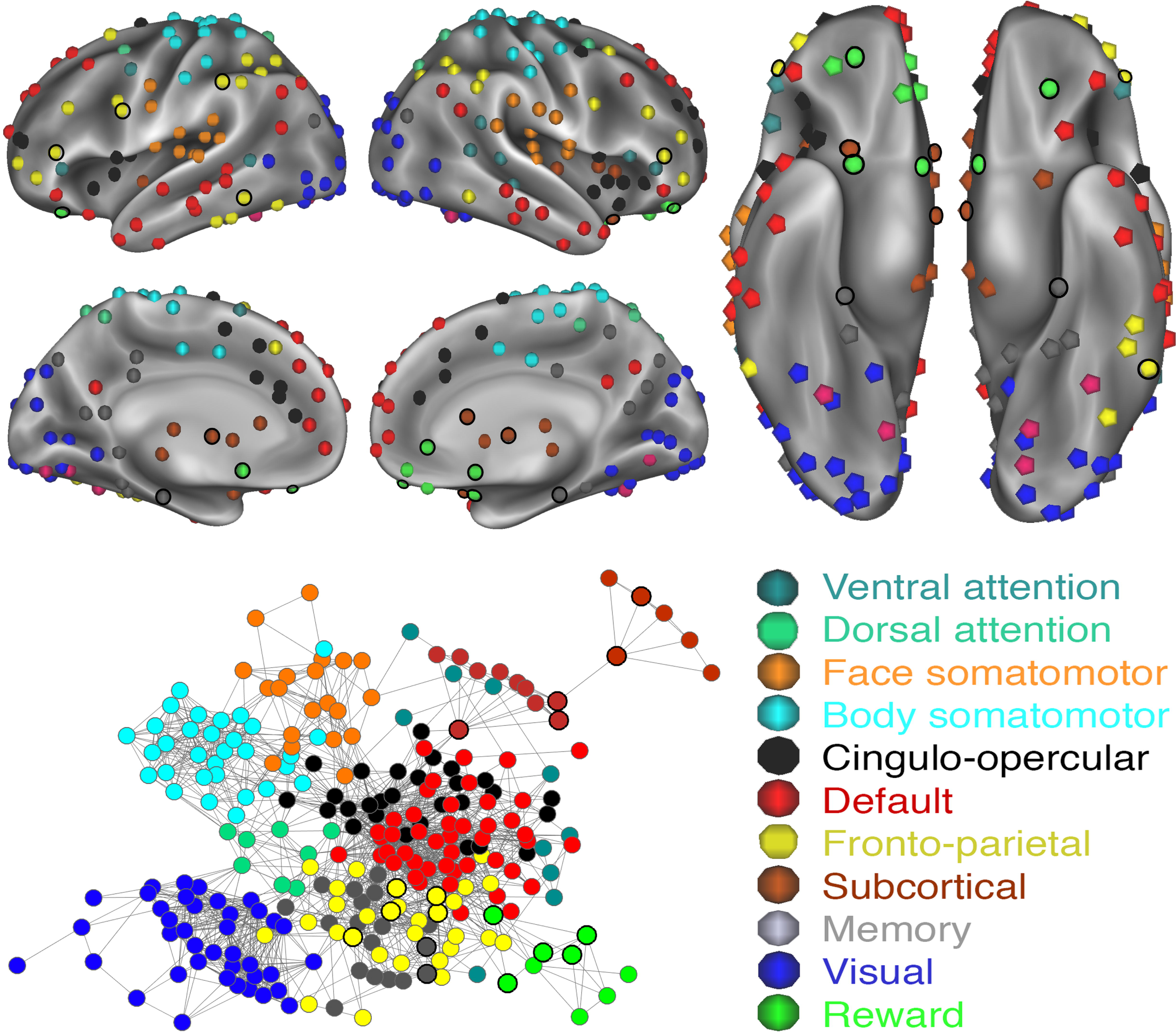
Community assignments displayed on an inflated cortical surface (top) at 5% tie density. A spring-embedded layout of the network graph visualizing network topology (bottom) displays nodes and community assignments at 5% tie density. Modified regions that were adapted from NAcc and OFC RSFC seed-based analyses are outlined in bold.

While the tie-density (defined above) is a useful way to identify the strongest percentage of connections and determine which connection should be submitted to a community assignment algorithm, it preferentially selects only the strongest connections, disregarding weak connections which have previously been shown to be underlie individual differences (Santarnecchi, Galli, Polizzotto, Rossi, & Rossi, 2014). In order to identify connections which are not necessarily the strongest connections but may be important we used a disparity filter algorithm to construct a multiscale backbone of the network (Serrano, Boguñá, & Vespignani, 2009). The term multiscale backbone simply means that connections across a variety of connectivity strengths which may be important for information transfer and maintaining the overall structure of the network optimally were included. Examples of where a multiscale backbone maintains relationships between network nodes better than simply using tie-density include complex networks such as the internet, food chain webs and the brain, all of which contain connections between nodes that are important across multiple orders of magnitude. The disparity filter algorithm identifies connections between nodes which significantly deviate from a null model of random local assignment of weights to edges thereby preserving much of the network structure while decreasing the total number of connections. In the current work, the resulting multiscale network backbone can then be used to identify connections which may be important for information transfer between a possible reward system and specific self-regulatory and control systems of interest.

### Visualization

All results were transformed into MNI space (Montreal Neurological Institute) and mapped onto the Conte69 mid-thickness surfaces or volume for visualization (Van Essen, Glasser, Dierker, Harwell, & Coalson, 2012). Group results were viewed in Connectome Workbench Version 1.1.1 (Marcus, Fotenos, Csernansky, Morris, & Buckner, 2010). Multiscale backbone results were visualized on inflated lateral, medial, and ventral cortical surfaces using BrainNet Viewer (Xia et al., 2013).

## Results

### Automated Segmentation of NAcc

Individual subject regions-of-interest (ROIs) for the left and right NAcc were defined using an automated segmentation algorithm (see Methods). The resulting NAcc masks were highly reliable across subjects with peak probabilities of 100% at [−8, 10, −10] and [9, 10, −11] (MNI coordinates). Fifty percent of the subjects had overlap in 900 and 880 voxels (mm^2^) for left and right NAcc masks, respectively. Seventy-five percent of the subjects shared overlap in 500 and 428 voxels (mm^2^). Mean volume of normalized individually defined subcortical regions are available in Table SI.

### Seed-based resting-state functional connectivity

Voxel-based RSFC maps were created for bilateral NAcc using the individually-defined masks from the automated segmentation and for OFC using spherical seeds based on peaks of NAcc connectivity which were proximal to regions identified in previous work (Wagner et al., 2012). NAcc demonstrated strong RSFC with the contralateral NAcc and bilateral regions of the medial and lateral OFC, thalamus, hippocampus, and midbrain/ventral tegmental area (Figure 1).

To determine if seed-based RSFC for these regions was similar to task-based activation patterns observed in studies of reward, bilateral NAcc and OFC seed maps were submitted to Neurosynth Decoder, an online tool to determine the similarity between the RSFC seed map and thousands of reverse-inference maps generated through automated meta-analyses of task-based fMRI studies (Yarkoni et al., 2011). Results of the reverse inference decoding analysis revealed the greatest similarity across all non-anatomical terms in the database between NAcc and OFC RSFC seed maps and the feature term “reward” (NAcc r = 0.58; OFC r = 0.27; Figure 1). Table I includes non-anatomical and anatomical terms which were showed the strongest correlation with NAcc (left) and OFC (right) RSFC seed maps.

### Community detection

Consistent with Power et al. (2011), communities were identified that included default, visual, fronto-parietal, cingulo-opercular/salience, attention, and somatomotor systems that were stable across a range of tie-densities (2%-20%). Additional systems from Power et al., (2011) were identified at lower tie-density thresholds including cerebellar (18%), subcortical (15%) and memory (8%) systems. The somatomotor system was further subdivided into and mouth and body somatomotor systems (6%). Figure 2 shows system assignments at a representative 5% tie-density.

In contrast to Power et al. (2011), however, the present work observed a distinct system that included the NAcc bilaterally and multiple regions of the medial and lateral OFC. This putative reward system was stable across a broad range of tie densities (4% to 17%). Two points are worth noting regarding this system. First, not all regions identified from NAcc and OFC RSFC seed maps converged into this reward system. At 5% tie-density, five of the regions identified in seed-based RSFC maps were assigned to the fronto-parietal system (lateral and dorsolateral frontal cortex), five were assigned to a subcortical system (left and right thalamus, left and right caudate, right putamen), two were assigned to a memory system (left and right hippocampus), and the VTA shared no connections to other nodes (at slightly higher tie-densities, e.g., 6% tie density, VTA was assigned to the memory system) (Figure 3; bold nodes). Second, four original nodes from the Power et al. (2011) graph were assigned to the “reward” system in the present analysis. In Power et al. (2011), these regions were previously assigned to fronto-parietal, default and an unlabeled system (Figure 3; green nodes without bold emphasis). At the highest tie-densities tested (18-20%) four of the nine reward nodes were subsumed by the default system (left and right NAcc and 0, 43, −7; 8, 48, −15), whereas the remaining five reward nodes were subsumed by the fronto-parietal system (23, 33, −13; −24, 35, −13; 8, 41, −24; 24, 45, −15; 34, 38, −12).

**Figure 3.**
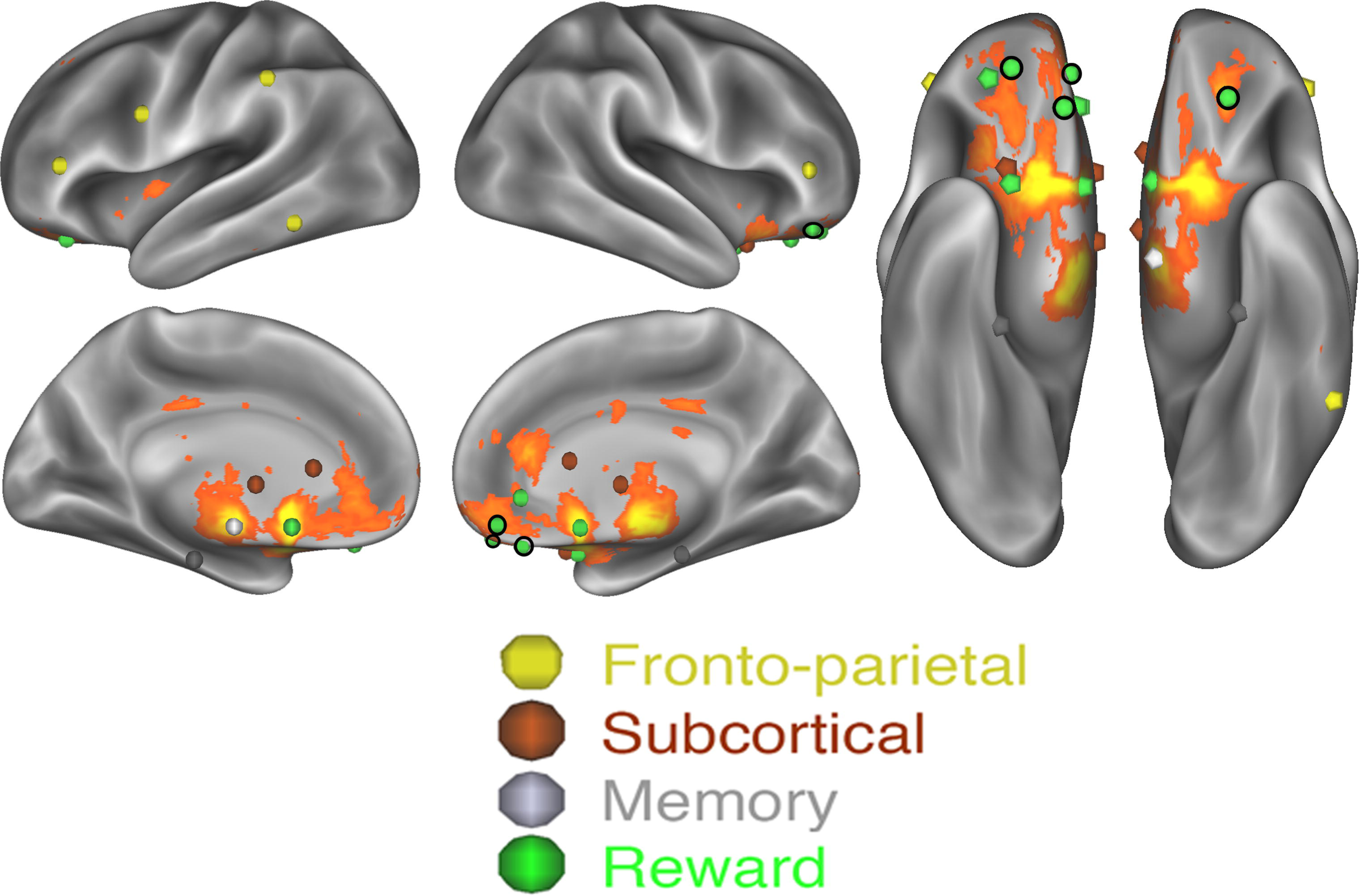
Candidate reward regions identified from seed-based RSFC (highlighted bold, any color) and reward system regions identified from graph analysis (green) are displayed on the task-based meta-analysis of reward (Yarkoni et al., 2011). Candidate regions are highlighted in bold of any color. Several candidate reward regions were assigned to other systems (non-green bold regions) and several additional nodes that were not directly identified in seed-based NAcc and OFC RSFC were assigned to the reward system via community detection (green non-bolded regions).

Within the reward system, stability was heterogeneous across regions; eight of the nine regions identified as being part of the reward system were stable between tie-densities of 4%-17% (five of the eight remained in this system down to 2%); at higher tie densities three of these regions were associated with the default network and five were associated with the fronto-parietal network. The ninth region, a medial OFC region (MNI: 0, 43, −7), was associated with the reward network across a more limited range of tie-densities (4%-8%), making it a less stable member of the reward network. This region was associated with the default network at tie-densities of 9% and higher. A tenth region was attached to the reward system at two tie-densities (6 - 7%), was part of the fronto-parietal at higher tie-densities and part of its own community at lower tie-densities. As such, this region should likely not be identified as part of the reward system.

To test the hypothesis that there is a reward system with preferentially coupled regions permutation testing was performed across randomly selected groups of 25 individuals. The null-hypothesis was that no independent system would be identified from the candidate regions. At the lowest tie-density (2%) five reward regions were identified as being clustered as a preferentially coupled system. Unsurprisingly, these are the same regions identified as being part of the reward system by the Infomap algorithm with all 828 individuals. At 4% tie-density these five regions were still significantly clustered with each other and also with the remaining four regions being part of the reward system in 757 or more of the 1000 permutations. A modified medial OFC region (MNI: 0, 43, −7) which was part of the reward system at five tie-densities was part of the reward system in 905/1000 permutations, not meeting our significance criteria. Simply, at 4% tie-density, eight of nine regions meet our criteria to reject the null-hypothesis and that they are preferentially coupled at p<=0.001. The medial OFC region (MNI: 0, 43, −7) does not meet these criteria and therefore we cannot say it is preferentially connected to the other reward regions in a significant manner but that it is coupled to those other regions more often than not. Critically, the roughly 11% of the time it was not part of the reward system, it was coupled with default system regions. At higher tie-densities the 8 members of the reward system identified above were preferentially coupled in a significantly manner (p<0.05) until 17% tie-density where, some of the regions dropped below p<0.05, which is unsurprising given that the N=828 group-level Infomap assignments are then subsumed by supraordinate systems (fronto-parietal and reward).

The overall spring-embedded network organization was similar to previously identified layouts as observed across large numbers of individuals, with control and attention systems such as the fronto-parietal, cingulo-opercular, ventral attention and dorsal attention systems being centrally located in the graph (Power et al., 2011). Canonical processing systems such as visual and somatomotor systems were located on the periphery of the graph. Across all tie-densities, the reward system was placed on the periphery of the graph. At low tie-densities (5%) the reward system connected with the default system (N=13 edges), cingulo-opercular (N=1) and subcortical (N=1) systems. At higher tie-densities (10%) the reward system was more densely interconnected with the default (N=34), fronto-parietal (N=11), cingulo-opercular (N=5), memory (N=3), dorsal attention (N=2) and subcortical (N=1) systems. As tie-densities increased further (> 17%), all reward system nodes were subsumed by default and fronto-parietal systems (Figure 4).

**Figure 4.**
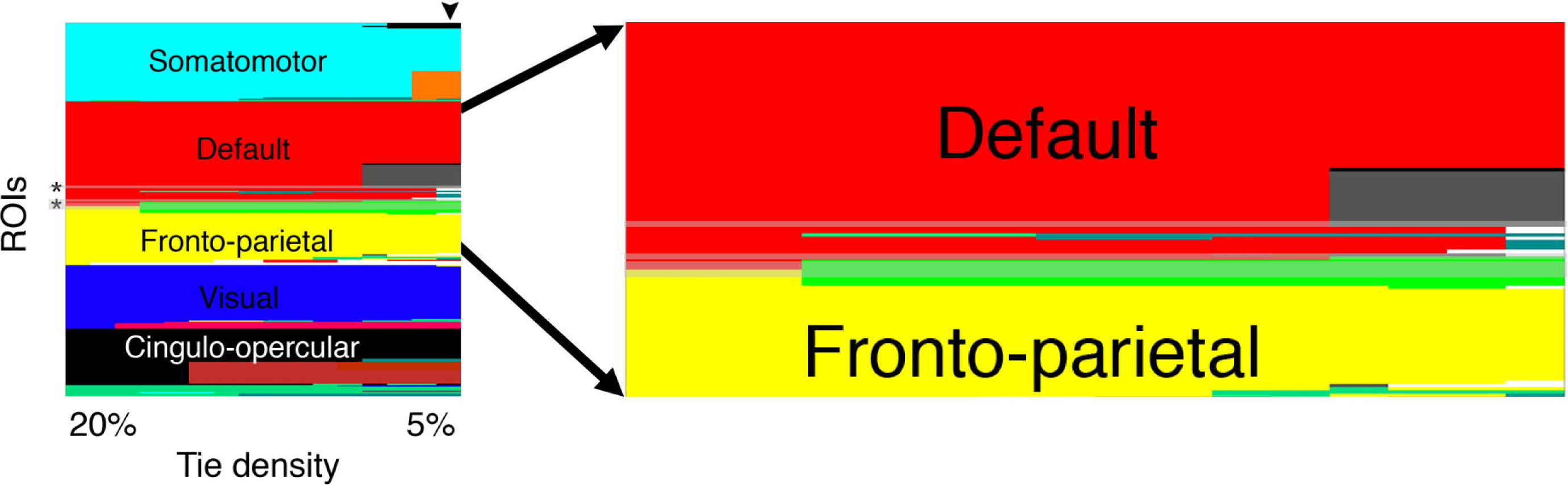
Plots are shown for node assignments into systems (left), with an enlarged view (right) which includes default, fronto-parietal, memory (in grey) and reward (in green) at tie densities from 20% down to 5% in 1% steps. Shaded rows and * represent modified ROIs.

Community detection is sensitive to the nodes which are included and the connectivity threshold (tie-density) at which it is performed. For a system to be considered stable it should be identified across a variety of tie-densities. To determine the consistency of community detection results across systems and tie-densities, a “system stability” metric was calculated which measures the relative amount of tie-densities that a community was identified for a given node. Superordinate systems, i.e. systems identified at 20% tie-density included default, fronto-parietal, visual, somatomotor, and cingulo-opercular/salience. Subordinate systems (i.e. systems which were identified only at tie-densities less than 20%) included reward, memory, subcortical, attention (dorsal and ventral), cerebellar, and lateral somatomotor systems. Critically, the newly identified reward system demonstrated system-wide consistency over a wide-range of tie-densities, and, as such, displayed high mean system stability relative to other subordinate systems (Table III). While community detection is sensitive to the nodes which are included and tie-density at which the analysis is performed, in the current work it was relatively stable, with qualitatively similar results under a variety of circumstances. Results across all analysis reliably found a reward system which contained similar nodes (more thoroughly described in Supplemental Material), including comparisons of high and low motion individuals, longest and shortest scan durations (10 vs 20 minutes).

**Table III.**
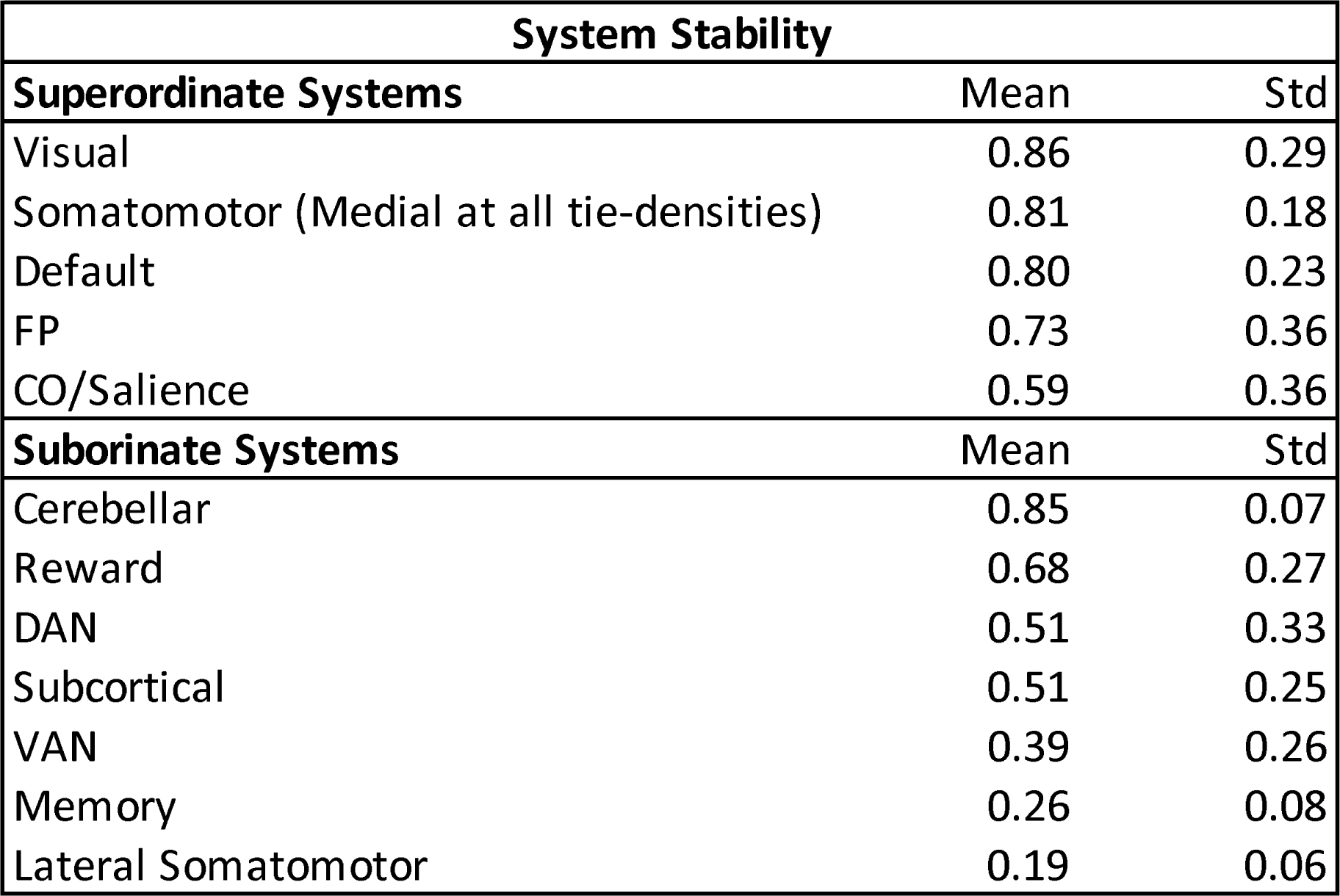
Stability of superordinate and subordinate systems across tie-densities. For each node that is a member of a given system at any tie-density between 2% and 20%, those nodes were selected, and the stability proportion metric was calculated. Stability proportion was calculated as the number of tie-densities a given system was observed at, divided by the number of tie-densities (19, 2%-20%). Systems with appearances in less than 1% of 271×19 matrix were excluded, including a second subcortical system.

An additional advantage of using graph-theory analyses on RSFC data is that it affords an opportunity to characterize points of interaction between systems. To identify points of interaction between the reward system and other systems, the entire network was submitted to a network reduction algorithm to identify the multiscale backbone of the system (Serrano et al., 2009). Backbone connectivity through reward system nodes was restricted to other reward system nodes and default system nodes (thresholded at p < 0.01; Figure 5). At this threshold, no other systems were identified as having strong connections to reward system nodes.

**Figure 5.**
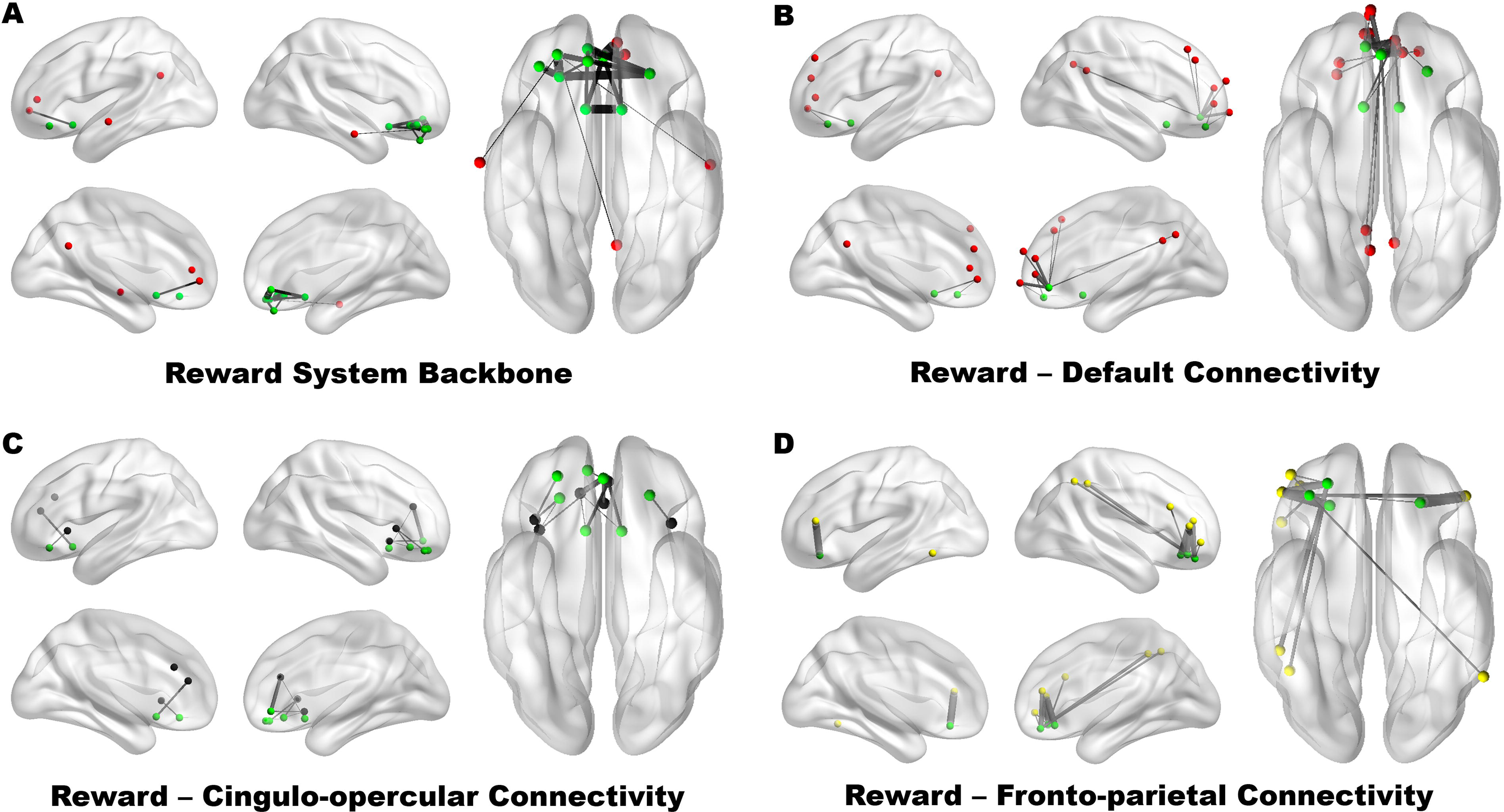
(A) Multiscale backbone connectivity (p < 0.01) through reward system nodes demonstrate strongest between-system integration with the default system. (B-D) The top 5% of connections between default (B, red), cingulo-opercular (C, black), and fronto-parietal control (D, yellow) and reward system nodes (green) reveal greater connectivity between frontal regions of default, cingulo-opercular, and fronto-parietal control systems and OFC regions of the reward system. Between-system connectivity with the NAcc was restricted to the dorsal anterior cingulate cortex (BA 32; cingulo-opercular system) and the medial prefrontal cortex (BA 10; default system). Intra-system connectivity is not shown. Regions and connections are displayed on inflated lateral, medial, and ventral cortical surfaces using BrainNet Viewer (Xia et al., 2013).

To identify potential interactions between the reward system and systems implicated in self-regulation (e.g., default, cingulo-opercular, and fronto-parietal systems) (Kelley et al., 2015), the top 5% of between-system connections from reward system regions to regions of these putative control systems were identified (Figure 5). Across all putative control systems, the majority of between-system connections were observed between OFC regions of the reward system and prefrontal regions of default, cingulo-opercular, and fronto-parietal systems. This was particularly true for between-system connectivity with the fronto-parietal control system, which was entirely driven by lateral OFC regions, which are often more associated with punishment and motor inhibition (Haber & Knutson, 2010). By contrast, the primary between-system connections for the NAcc were with the dorsal anterior cingulate cortex extending into the middle frontal gyrus (BA 32; cingulo-opercular system) and the medial prefrontal cortex (BA 10; default system) for reward to cingulo-opercular and reward to default system connectivity.

## Discussion

The current study applied seed and graph-theory based approaches to RSFC data from a large sample of healthy individuals using a set of nodes modified from previous work (Power et al., 2011). Seed-based RSFC from canonical reward regions demonstrated a high degree of similarity with task-based meta-analyses on the feature term reward. Peaks of connectivity from these putative reward regions were identified and used to modify an existing set of nodes (Power et al., 2011). Three possible outcomes were considered: 1) no resting-state reward system (i.e., all of the candidate nodes are members of other RSFC systems); 2) all of the candidate reward regions clustered into a distinct system, and 3) a subset of the regions comprised a distinct system. Graph theory analyses that included this modified set of regions identified a subset which were preferentially coupled and formed a distinct community across a wide variety of tie-densities, falling into the third possible outcome. Permutation testing suggests that at eight out of the nine regions in this system are preferentially coupled more than 95% of the time. Perhaps unsurprising, this distinct “reward” community or system shows strong overlap with regions identified through task-based analyses of reward. Although seed-based and graph-based reward maps were consistent in some respects, notable differences were observed across the two approaches. We consider each in turn.

### Seed-based RSFC show robust overlap with task-based reward activity

Seed-based RSFC from individually-defined NAcc regions showed robust overlap with task-based studies of reward. Peak correlations were observed in the striatum, medial and lateral OFC, and the hippocampus. These findings are consistent with prior studies of NAcc RSFC using spherical regions-of-interest (Di Martino et al., 2008). However, the present work also identified RSFC peaks in regions not previously reported by Di Martino and colleagues including thalamus and midbrain/ventral tegmental area. The extent to which the present findings identify minor differences from prior work may reflect differences owing to a different cohort of subjects, increased sample size, the greater specificity afforded by using individual NAcc segmentations instead of spherical nodes and different preprocessing approaches.

Seed-based RSFC for the lateral OFC showed strong overlap with RSFC for NAcc. The OFC was strongly correlated with lateral regions of the left and right OFC, the medial OFC, and bilateral NAcc. Additional connectivity was observed in lateral and dorsolateral frontal regions as well as lateral parietal regions that were not present in the NAcc RSFC map. These regions were identified in subsequent community assignment analyses to belong to the fronto-parietal system, however— a finding that suggests that one potential point of communication between the reward system and other systems important for self-regulation and control may occur via interactions between dorsolateral prefrontal cortex and the lateral OFC (Figure 3). By contrast, the NAcc was more strongly correlated with regions of the default system (i.e., medial prefrontal cortex, inferior parietal lobule, and superior temporal gyrus), the memory system (i.e., parahippocampal gyrus) and subcortical structures (i.e. VTA/midbrain and thalamus).

Although the relationship between task-driven functional correlations and RSFC has been highlighted across the extant neuroimaging literature (Crossley et al., 2013; Smith et al., 2009; Tomasi, Wang, Wang, & Volkow, 2013), the exact nature of this relationship remains somewhat speculative. One possibility is that RSFC represents long-term Hebbian learning, such that populations of neurons across brain regions that fire together frequently strengthen spontaneous resting state correlations over time. As such, RSFC may represent the statistical history of co-activation patterns accumulated across the lifespan. In the present study, NAcc RSFC, and to a lesser extent OFC RSFC shared strong overlap with meta-analyses of task-based reward activity. Future studies may be able to capitalize on the similarity between RSFC and task-based reward patterns of activation to explore the development and interaction of the reward system with control systems, which have been argued to mature along different developmental trajectories (BJ J Casey, Jones, & Hare, 2008; Somerville & Casey, 2010) and may be influenced by genetic factors that influence the size and responsivity of reward system structures (Rapuano et al., 2017).

### Graph-theory identification of a reward system

The modified set of brain regions was submitted to a community detection algorithm where we expected one of three possible outcomes: 1) no resting-state reward system (i.e., all of the candidate nodes are members of other RSFC systems); 2) all of the candidate reward regions clustered into a distinct system, and 3) a subset of the regions comprised a distinct system. The first possibility was that there was no reward region, such that all of the reward-related nodes would disperse into several systems, but we did not observe this. The second possibility would be that all of these nodes were part of a larger, supraordinate system across most tie-densities observed. This was partially observed, in so much that several of the nodes were part of the supraordinate default system at extremely high tie-densities (18%+) and there was only a total of six systems observed. At lower tie-densities several (eight or nine depending on threshold) regions were clustered into a subordinate reward system, suggesting that the third possible outcome is what we are observing in the current study. We observe that several canonical reward regions form a preferentially coupled resting-state system; system membership included bilateral NAcc, the medial and lateral OFC, and ventromedial prefrontal cortex. This system was stable across a wide range of tie-densities (4-17%) with eight of the nine regions consistently assigned to the system across this range, showing a higher system stability score than multiple other well-established subordinate systems. Permutation testing results showed that these eight regions were significantly clustered in a preferential manner across multiple tie-densities, preferring to couple with each other than other systems, showing that the results mentioned here are very stable across individuals. The ninth region (medial OFC, MNI: 0, 43, −7) likely plays a role in reward-related activity as the top non-anatomical term for this region in Neurosynth is “default” followed by “reward”, but it does not meet our criteria for significance through permutation testing. It should be noted that qualitatively similar systems incorporating some of the ventromedial PFC and ventral striatum have been observed using different methods (Choi et al., 2012; Gerraty, Davidow, Wimmer, Kahn, & Shohamy, 2014). These complementary approaches suggest stability not just across tie-densities thresholds, but across datasets and methods. Furthermore, within our own dataset, we observe the reward system across multiple analyses. Individuals with high or low motion (mean frame-displacement or FD), longer or shorter scans (10 or 20 minutes) and even using slightly different node selections. Researchers sometimes use an exclusion distance of 20mm to make sure that proximity lead to artificial local systems being identified. The results presented in figures here do not include an exclusion distance, but nearly identical results are observed when this is done (see Supplemental Material for more information). Moreover, similar results are observed with weighted or binarized thresholded networks for each analysis.

In the current work we strove to optimize the chances of identifying a reward system which was preferentially connected at rest, through which we used a moderately intensive procedure to select nodes which would give us the best chance of identifying this system. After completing this, we backtracked and added bilateral nucleus accumbens to the original Power 264 region set to see if it would result in a preferentially coupled reward system (Table S2). In this is the case, a system which primarily contained OFC regions and NAcc was present from 2-18% tie-density. At 5% tie-density these 12 regions were identified as part of this system. It is likely not as specific as that identified using our ROI modified procedure as in addition to including bilateral NAcc and OFC regions it also included a cerebellar seed and a parietal seed. This further highlights the necessity for seed selection, particularly when attempting to optimize for a specific task such as attempting to identify a reward system.

With the exception of regions that reorganized into the reward system identified here, there was strong overlap in community assignments across studies. Similar to Power et al. (2011), cognitive control systems such as the frontal-parietal and cingulo-opercular systems were centrally placed within the graph whereas somatosensory-motor and visual systems were placed on the periphery of the graph. The reward system, as defined in the current analysis, was placed at the periphery of the graph and showed the most connectedness with default, cingulo-opercular, and fronto-parietal systems (at liberal tie-densities). The relative locations of subsystems within a graph provide insights into whether a system may play a more regulatory role (centrally located within the graph), by virtue of their stronger connectedness with multiple systems, or whether a system functions primarily as a processing system (located at the periphery of the graph) with strong within-system connectivity and relatively weaker between-system connectivity. In this context, the reward system is more similar to other, well-established processing systems, and, as such, is likely a target of self-regulatory control via interactions with systems more centrally placed within the graph. Indeed, recent work by Power et al., (2013) has identified cortical hubs in RSFC data that demonstrate strong between-system connectivity in a way that positions hubs to interact with multiple systems, and lesions to these hubs have been shown to result in profound behavioral impairments across a range of tasks (Warren et al., 2014).

The present results suggest that the majority of between-system connections to the reward system occur between putative control systems (default, cingulo-opercular, and fronto-parietal systems) and OFC regions of the reward system. Interestingly, the OFC seed region identified from task-based studies of reward shares its primary between-system connections with the fronto-parietal system, whereas the NAcc shares its primary between-system connections with a single region of the cingulo-opercular system (dorsal anterior cingulate cortex; BA 32) and a single region of the default system (MPFC; BA 10). Indeed, at liberal tie-densities (> 17%), the OFC seed region associates with the fronto-parietal system, and the NAcc associates with the default system. The MPFC region of the default system has been implicated in the representation of self and self-affect (Kelley et al., 2002; Moran, Macrae, Heatherton, Wyland, & Kelley, 2006), and recent work by Chavez and Heatherton (2016) has demonstrated that the structural integrity of white matter pathways linking NAcc to this region of MPFC is associated with individual differences in self-esteem. The differential connectivity patterns between OFC, NAcc and putative control systems may offer critical insights into the differing roles of these regions in motivating and evaluating appetitive behaviors that have so far been elusive given the common coactivation of these regions during task-based studies of reward. Although the current results do not demonstrate direct evidence of regulation paths between control and reward networks, the present work identifies potential points of interaction with the multiscale network backbone analysis. Future work is needed to clarify the extent to which these connections may play a role in enhancing or regulating appetitive cravings and behavior. Towards that end, a better understanding of the cortical architecture that permits regulation of these reward regions may ultimately permit more targeted intervention strategies for the treatment of maladaptive habits and addictions.

The most notable difference between Power et al. (2011) and the present study was the inclusion of NAcc nodes in the present study and the absence of NAcc nodes in the prior work. Previous RSFC work using a winner-take-all method using only subcortical regions associated much of the ventral striatum with a limbic system that included portions of the temporal poles (Choi et al., 2012). Davis and colleagues (2013) demonstrated that NAcc system membership may depend in part on individual differences in reward impulsivity. In a high impulsivity group, NAcc clustered with other subcortical regions such as caudate, amygdala, hippocampus, thalamus and brainstem whereas in a low impulsivity group, subcortical regions clustered with top-down cognitive control regions, such as medial and lateral portions of the prefrontal cortex.

Importantly, however, not all of the candidate reward system regions identified from seed-based RSFC analyses were formally assigned to the reward system identified using graph-based network analyses, highlighting the complementary nature of the two approaches (Figure 3). Two important distinctions can be made. First, there is near complete overlap between regions assigned to the reward system in the graph analysis and the meta-analysis of task-based studies of reward, even though some of these regions (e.g., anterior OFC nodes) were not identified as peaks in the seed-based NAcc and OFC RSFC maps. Second, some of the candidate “reward” regions identified from the seed-based RSFC maps were more strongly connected to other systems in the graph analysis (e.g., inferior frontal gyrus, hippocampus, and thalamus), even though they overlapped with commonly activated regions in task-based studies of reward. This latter finding suggests that task-based studies of reward engage multiple systems (reward, subcortical, memory, and fronto-parietal systems) and that seed-based RSFC and task-based meta-analyses may be less well-suited to capture these distinctions. As such, it is not sufficient to propose a reward system based on voxel-based seed maps, nor is it sufficient to do so using meta-analyses of task-based reward studies. By identifying networks which are consistent across a variety of methods, we can better characterize these networks refine them as needed.

Graph-theoretical approaches are better suited to segregate brain regions into brain systems, however this approach too is limited as community assignments are highly dependent on how nodes are defined and on the correlation thresholding applied to the graph. The present study used nearly identical methods from prior work (Power et al., 2014) on a set of brain regions modified from Power and colleagues (2011) and identified a reward system not previously observed using the original set of regions. The regions were purposefully modified using seed-based RSFC to refine the graph to include regions commonly associated with reward processing and their neighbors, and, when possible, regions were further refined on a subject-by-subject basis using anatomical segmentation. Similar modifications of the canonical Power et al. (2011) graph were previously used to demonstrate distinct neurophysiological subtypes of depression using RSFC data (Drysdale et al., 2016).

### Future Directions and Limitations

We do not suggest that the current set of regions which constitute the reward system as we have defined it is necessarily the optimal one, but instead a starting place from which to expand upon. With this in mind, the reward system delineated in the current work defines a set of regions that future studies may capitalize on to better understand reward motivation, appetitive behavior and self-regulatory control. Other groups have identified a similar system or set of brain regions using other methods, without the goal of specifically defining a reward system (Choi et al., 2012; Davis et al., 2013). The overlap between the current results and previous work, particularly across methods suggests the concept of a set of brain regions which support reward processing and are highly coupled at rest. This also suggests, as one might expect, that the regions in this preferentially coupled system subserve multiple purposes. Although additional brain regions are often co-activated with these core regions during reward processing, it would be inappropriate to include them in the definition of a reward system as they are better members of other RSFC brain systems. The current set of regions is by no means comprehensive, but instead a target approach to modify the Power 264 region set to be focused towards reward. There are many other regions which could possibly be included which play a role in reward processing, such as substantia nigra and amygdala, but neither of those showed up in our connectivity-based peak selection process. Furthermore, neither the amygdala and substantia nigra were part of the reward system when they were added in a secondary analysis. Over the past decade there has been a growing interest in using RSFC data to parcellate the human cortex into cortical areas. Future work may be able to leverage these more sophisticated parcellation schemes (e.g., Glasser et al., 2016; Gordon et al., 2014) to further refine reward system identification and characterization. Given enough data on an individual, subject-specific brain parcellation may also further this endeavor and could lead to an increased sensitivity to detecting individual differences in reward-related behavior (Gordon, Laumann, Adeyemo, & Petersen, 2015; Laumann et al., 2015). Methods to increase functional alignment across cortex in individuals such as task-based hyperalignment (Guntupalli et al., 2016) or connectivity based hyperalignment (Guntupalli & Haxby, 2017) would increase alignment across individuals and may further help delineate the boundaries of the reward system across individuals. Future work could focus on optimizing the integration of cortical surface parcellations with subcortical time-series, something that could provide promise for individual parcellations or hyperalignment methods. The current work, with continued optimization of a reward-focused parcellation or hyperalignment methods which integrate subcortical regions will provide tools to more accurately compare individual and group differences as some are starting to do (Ma et al., preprint: 10.1101/296012). A critical future test of an “optimized” reward system would be increased utility in identifying individual differences in reward behavior. In parallel with this, regions responsible for self-regulation and inhibition should continue to be interrogated, perhaps within the framework of the connections identified in the current work through the multiscale network backbone. Functional gradients are known to exist within the ventral striatum and the OFC. Methods to identify and characterize these gradients in RSFC, somewhat similarly to methods in Barnes et al. (2010) and Choi et al. (2010), mapping maximal connectivity from subcortical regions to maximal connectivity in OFC/prefrontal cortex may help determine the extent to which these gradients exist. Ideally, an “optimized” reward system definition combined with knowledge of these gradients would help researchers elucidate individual differences in reward and self-regulatory behaviors.

## Summary

The current study characterized the systems-level organization of putative reward regions of at rest, identifying a heterogeneous set of regions including cortical and subcortical nodes which are preferentially coupled to each other and that may reflect a resting reward system. The high degree of similarity between seed-based RSFC maps of putative reward regions and meta-analytical maps of task-based reward suggested a putative reward system that was formally tested using graph-theory analysis. To afford the best chance of identifying a reward system, a previously published set of nodes (Power et al., 2011) was modified to include reward-sensitive brain regions and their neighbors. We then used a data-driven community assignment approach and identified a reliable reward system consisting of regions that both overlapped with and constrained task-based reward activation maps. By delineating and characterizing the reward system at rest in this way, future work may benefit from more spatially constrained analyses that target individual and group differences in reward processing and self-control. Thus far, the regulation of these reward regions has been a highly debated topic (Kelley et al., 2015), in part because the reward system has not been formally defined despite a consistent pattern of task-based reward activity in the literature and also because it is difficult to identify a commonly activated set of brain regions associated with self-regulation. Although it is not yet clear whether RSFC can be used to assess directionality of information flow between regions and systems, it can be used to constrain the brain regions that comprise a functionally-connected reward system and to identify critical through-points between reward and putative control systems that can be targeted in future intervention studies.

## Acknowledgements

This work was supported by NIH DA022582, MH059282, AA021347, HL114092, NSF BCS-0746220, the Neukom Institute for Computational Science at Dartmouth College, and the Dartmouth Brain Imaging Center. We would like to especially thank the Editors and three anonymous Reviewers for their valuable feedback.

